# *In vivo* fate of systemically administered encapsulin protein nanocages and implications for their use in targeted drug delivery

**DOI:** 10.1101/2023.07.16.549228

**Authors:** Claire Rennie, Caitlin Sives, India Boyton, Dennis Diaz, Catherine A Gorrie, Orazio Vittorio, Lyndsey Collins-Praino, Andrew Care

## Abstract

Encapsulins, self-assembling protein nanocages derived from prokaryotes, are promising nanoparticle-based drug delivery systems (NDDS). However, the *in vivo* behavior and fate of encapsulins are poorly understood. In this pre-clinical study, we probe the interactions between the model encapsulin from *Thermotoga maritima* (TmEnc) and key biological barriers encountered by NDDS. Here, a purified TmEnc formulation that exhibited colloidal stability, storability, and blood compatibility was intravenously injected into BALB/c mice. TmEnc had an excellent nanosafety profile, with no abnormal weight loss or gross pathology observed, and only temporary alterations in toxicity biomarkers detected. Notably, TmEnc demonstrated immunogenic properties, inducing the generation of nanocage-specific IgM and IgG antibodies, but without any prolonged pro-inflammatory effects. An absence of antibody cross-reactivity also suggested immune-orthogonality among encapsulins systems. Moreover, TmEnc formed a serum-derived protein corona on its surface which changed dynamically and appeared to play a role in immune recognition. TmEnc’s biodistribution profile further revealed its sequestration from the blood circulation by the liver and then biodegraded within Kupffer cells, thus indicating clearance via the mononuclear phagocyte system. Collectively, these findings provide critical insights into how encapsulins behave *in vivo,* thereby informing their future design, modification, and application in targeted drug delivery.

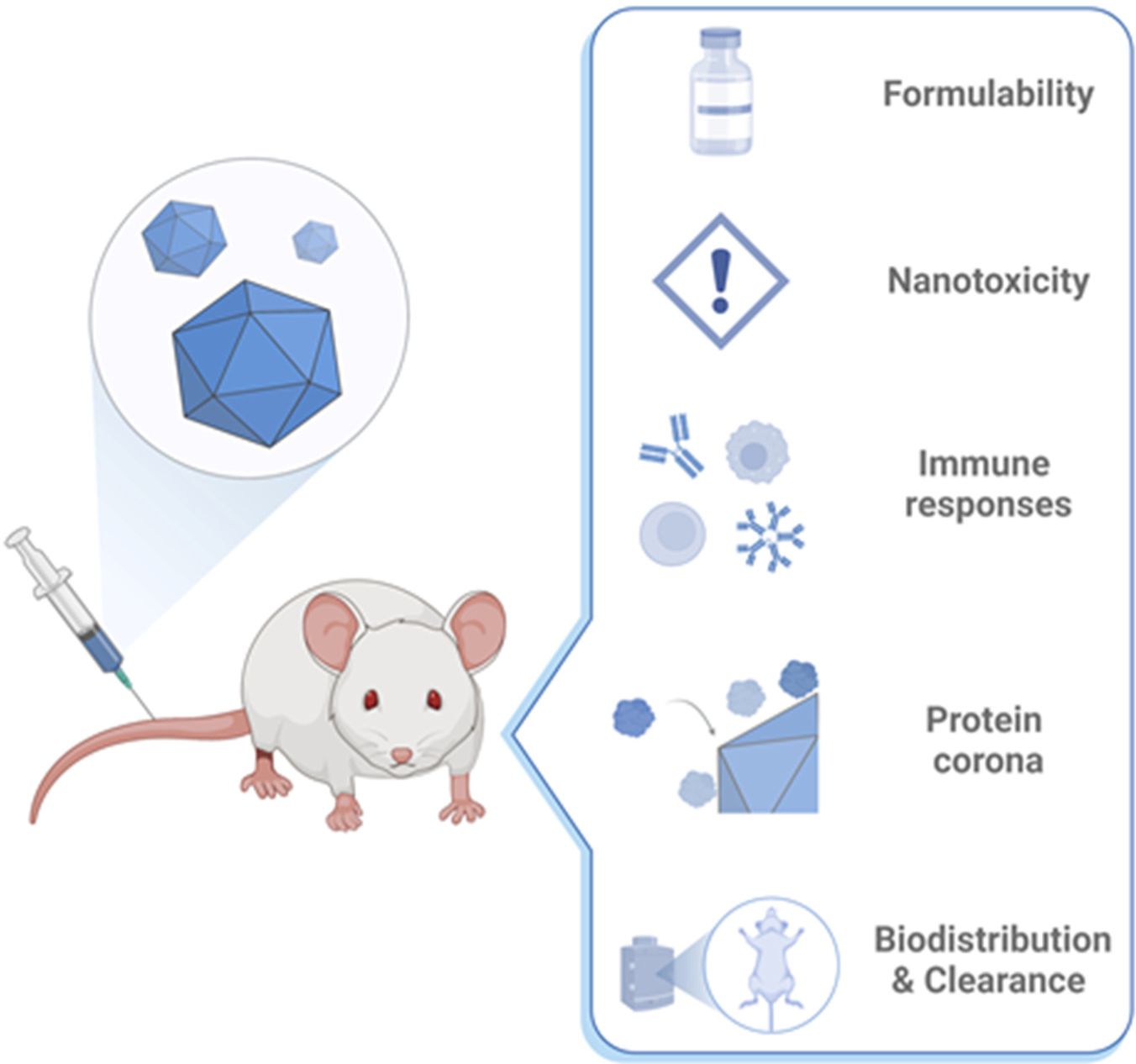

## 1. Introduction

Protein nanocages (PNCs) self-assemble from multiple protein subunits into highly-organized macromolecular structures.[1] They can be derived from a multitude of natural sources (e.g., viral capsids, ferritins, heat shock proteins, chaperonins, bacterial compartments), or in some cases, *de novo* designed.[1–3] PNCs have inherent features that make them attractive nanoparticle-based drug delivery systems (NDDSs).[3, 4] This includes hollow interior cavities for drug encapsulation; exterior surfaces to display disease-targeting ligands; and protein subunit interfaces that enable controlled drug release. In contrast to conventional synthetic nanoparticles, the structural and functional properties of PNCs can be both genetically and chemically manipulated with high precision. Furthermore, they are stable in physiological fluids, non-toxic, biodegradable, and offer reliable manufacturing due to their biological synthesis.[4] As a result, an array of custom-engineered PNCs have been developed for the targeted delivery of therapeutics (e.g., nucleic acids, proteins, small molecule drugs) for the treatment of various diseases, primarily cancer.[4]

Encapsulins are a newly established class of pseudo-organelles found inside many prokaryotes. They self-assemble from identical protein subunits into semipermeable nanocages that exhibit icosahedral symmetries: *T* = 1 (60-mer, 20−24 nm), *T* = 3 (180-mer, 30−32 nm), or *T* = 4 (240-mer, 43 nm).[5, 6] A unique feature of encapsulins is their ability to selectively encase native cargo proteins (e.g., enzymes) tagged with a short encapsulation signal peptide (ESig), a mechanism that can be readily co-opted to load the PNCs with foreign cargo.[7] Given their structural and functional modularity, encapsulins have attracted increasing interest as versatile platforms for biocatalysis, bionanotechnology, and biomedicine.[6–8]

The encapsulin derived from the bacterium *Thermotoga maritima* (TmEnc: *T* = 1) is the most extensively studied.[9] TmEnc is widely used as a model system to prototype encapsulin engineering for different practical applications.[10–21] Toward targeted drug delivery, the Kang lab modified TmEnc to display cancer-targeting peptides and the acid-sensitive chemotherapy pro-drug aldoxorubicin (AlDox) on its outer surface.[14] The drug-coated encapsulin was shown to selectively enter liver cancer cells, where AlDox was released and activated in acidic lysosomal compartments, leading to intracellular delivery and tumor cell death. In a different approach, we recently adapted encapsulins’ cargo loading mechanism for therapeutic protein delivery. Here, TmEnc was loaded with the ESig-tagged protein photosensitizer mini-Singlet Oxygen Generator (mSOG), forming an mSOG-loaded encapsulin (TmEnc-mSOG) nanoreactor that generated reactive oxygen species (ROS) under blue-light irradiation. After passive delivery into lung cancer cells, light-triggered TmEnc-mSOG produced intracellular ROS that induced oxidative stress and reduced cell viability, thus demonstrating photodynamic therapy (PDT).[13] Others later directly fused an antibody mimic (i.e., DARPin) to the exterior surface of the TmEnc-mSOG nanoreactor, enabling targeted PDT of breast cancer cells.[22]

While it is evident that encapsulins are exciting prospective NDDSs, TmEnc’s capacity for cell-specific targeting and drug delivery has only been validated to date using *in vitro* models.[10, 12–14, 19, 21, 22] In fact, *in vivo* studies of TmEnc have primarily focused on its use as an antigen carrier in proof-of-concept vaccine development. Examples include prophylactic/therapeutic vaccines designed against viruses (i.e., Epstein-Barr, Influenza, HIV,) as well as cancer.[11, 23–25]. Unlike vaccines, which target the immune system and are most often administered intramuscularly or subcutaneously, an NDDS is typically given *via* intravenous (IV) injection. This administration route offers direct entry into the bloodstream which serves as a distribution network that enables an NDDS to access target organs and tissues throughout the body (e.g., tumors).[26] However, an NDDS must overcome a distinct set of physiological and biological barriers in order to reach its intended site-of-action, such as immune system clearance, off-target accumulation in healthy tissues, and non-specific uptake by normal cells.[26–28] For this reason, understanding the *in vivo* behavior and fate of systemically administered TmEnc is a vital prerequisite for its development into a viable NDDS.

In this pre-clinical study, we have elucidated, for the first-time, the formulability, nanosafety, immunogenicity, protein corona, biodistribution, and clearance of IV injected TmEnc into healthy BALB/c mice. Our results provide new and critical insights into how encapsulins behave *in vivo,* thus informing their future design, modification, and application in targeted drug delivery.

## 2. Results and discussion

### Protein nanocage production, purification and characterization

To ensure safe and effective clinical utilization, high-quality NDDS formulations should be consistent, reproducible, stable, and storable.[26, 29] Unlike synthetic nanoparticles, PNCs are synthesized by biological systems thus enabling streamlined biomanufacturing processes with fewer production steps and minimal inter-batch variability.[2, 30] In this study, TmEnc nanocages were recombinantly produced in *Escherichia coli* (*E. coli*) and purified by size-exclusion (SEC) and anion-exchange (AEX) chromatography (**Figure S1**), and then biophysically characterized (**Figure 1**).[5] SDS-PAGE confirmed the purification of TmEnc (TmEnc_subunit_; 30.5 kDa) with densitometric gel analysis determining >90% purity (**Figure 1a**). Transmission electron microscopy (TEM) images of negatively stained samples visualized the correct formation of TmEnc into hollow spherical nanocages that were uniform in size and shape (**Figure 1b**). Dynamic light scattering (DLS) measurement of TmEnc revealed a mean hydrodynamic diameter of 20.9 ± 5.7 nm with a narrow size distribution (PDI < 0.2) (**Figure 1c**); this size is consistent with TmEnc’s reported crystal structure (*T* = 1; PBD: 3DKT).[9] In addition, zeta potential analysis further determined an overall negative surface charge of −8.4 mV for TmEnc. Together, this data showed that purified TmEnc nanocages were monodisperse and colloidally stable, which are important attributes for nanopharmaceutical formulations.

**Figure 1.**
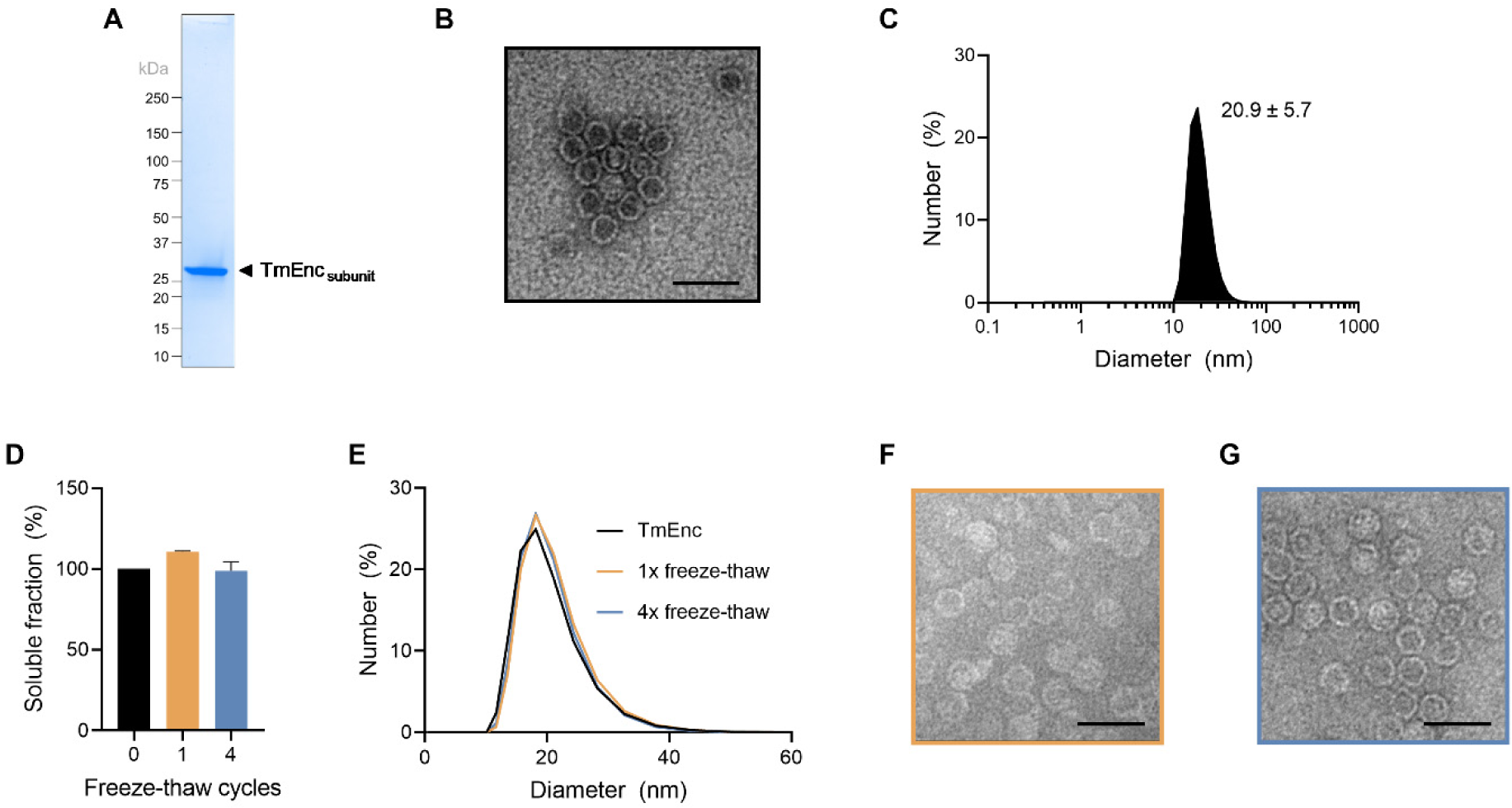
Production, biophysical characterization, and storage stability of TmEnc nanocages. **(a)** SDS-PAGE visualization of TmEnc nanocage (TmEnc_subunit_; 30.5 kDa) purified by sequential SEC and AEX. **(b)** TEM image showing formation of TmEnc into spherical nanocages. Scale bar = 50 nm. **(c)** DLS analysis of TmEnc determined a mean diameter of 20.9 ± 5.7 nm **(d)** SDS-PAGE densitometric quantification of soluble TmEnc recovered from purified samples exposed to 0x (black), 1x (yellow), or 4x (blue) freeze-thaw cycles at −80°C (see **Figure S2** for corresponding gel image). Results presented as mean ± SEM, *n* = 3. Both **(e)** DLS and **(f-g)** TEM images verified that the nanocage retained its stability and macrostructure after 1x (yellow) and 4x (blue) freeze-thaw cycles. Scale bars = 50 nm.

TmEnc’s protein shell displays exceptional resilience against extreme pH, high temperatures, chemical denaturation, and proteolytic degradation.[5, 18, 31] Nevertheless, protein cage formulations can become susceptible to structural disruption during storage, limiting their shelf-life.[32] To therefore assess storage stability, TmEnc nanocages were subjected to 1x or 4x freeze-thaw cycles at −80°C. SDS-PAGE densitometric analysis indicated that ≥ 99% of TmEnc remained soluble after multiple rounds of freeze-thawing (**Figure 1d and S2**), while DLS (**Figure 1e**) and TEM (**Figure 1f-g**) verified the preservation of the nanocage’s assembled macrostructure. Encouraged by the observed purity, stability, and storability of our TmEnc formulation, we proceeded to investigate the *in vivo* behavior of the nanocage.

### Blood compatibility and *in vivo* safety profile

IV injection is a clinically-relevant administration route for NDDSs which facilitates rapid distribution throughout the blood circulation system, and thus enables NDDS localisation within specific areas in the body. Up until now, pre-clinical investigations involving TmEnc have focused on its utility as an antigen-delivery system, primarily for vaccine development.[11, 24, 25] In such studies, nanocages are administered into animal models *via* subcutaneous or intramuscular injection to target draining lymph nodes and trigger humoral and cellular immune responses. With targeted drug delivery in mind, we therefore elected to intravenously administer TmEnc nanocages in all of our *in vivo* studies.

To ensure purified TmEnc was safe for IV administration, its blood compatibility was assessed *via* an *ex vivo* hemolysis assay (**Figure 2a**). Herein, isolated mouse red blood cells (RBCs) were incubated with varying concentrations of nanocages (50, 100, 200, and 400 µg/mL) at 37°C for 1 h. Upon completion, hemoglobin released by any lysed RBCs was determined by measuring absorbance at 400 nm.[33] At all tested concentrations, TmEnc induced less than 10% RBC lysis, which is well below the threshold (>25%) for hemolysis risk [34] and is consistent with the blood compatibility of other PNCs. [35–37] For example, Bruckman et al. found that tobacco mosaic virus (TMV) capsid exhibits non-hemolytic properties at similar concentrations. [36] Notably, the authors also highlighted that the ratio of TMV to RBCs in their *ex vivo* hemolysis assay was 1000-fold higher than what was subsequently delivered *in vivo*. According to this information, TmEnc nanocages are nonhemolytic and suitable for IV injection.

**Figure 2.**
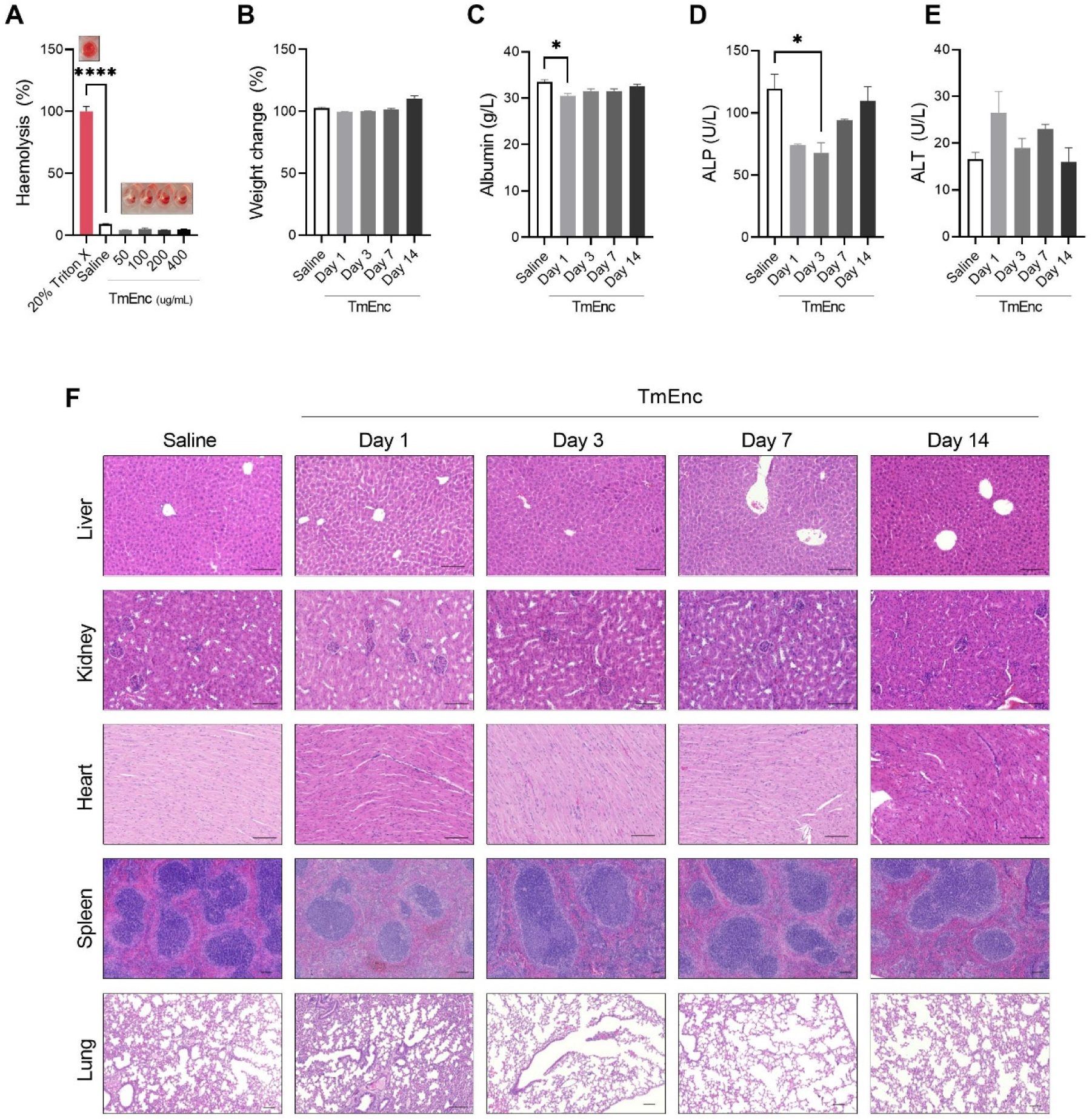
Safety profile of TmEnc nanocages. **(a)** *Ex vivo* haemolysis assay showing that TmEnc (0-400 µg/mL) did not lyse mouse red blood cells (inset: representative images of assay samples in microplate wells). TmEnc (5 mg/kg) was intravenously administered into BALB/c mice, which were euthanized 1-, 3-, 7- or 14-days after treatment with tissues and blood collected for analysis: **(b)** TmEnc did not cause any gross weight loss. Serum analysis indicated short-term changes in liver function markers **(c)** Serum albumin; **(d)** Alkaline phosphatase (ALP); and **(e)** Alanine aminotransferase (ALT). **(f)** Histological evaluation of organs (liver, kidney, heart, spleen, lung) by H&E staining at 1-, 3-, 7-, and 14-days post treatment demonstrated that TmEnc administration did not cause any abnormal gross pathology; 2.5X or 5X magnification; Scale bars = 100 µm. Results presented as mean ± SEM, one-way ANOVA, Dunnett’s, *n* = 4 (a and b) or Tukey’s, *n* = 2 (c-e) (*P<0.05, ***P<0.001, ****P<0.0001).

While there have been no reports of TmEnc inducing any observable nanotoxicity in pre-clinical animal studies,[11, 24, 25] a comprehensive *in vivo* safety profile is still lacking. In pursuit of this critical information, TmEnc (5 mg/kg) was systemically administered into healthy BALB/c mice *via* tail-vein injection (IV). Mouse groups (*n* = 4) were then euthanized at 1, 3, 7, or 14 days post-injection, with tissues and blood collected and subsequently analyzed using a battery of hematological, biochemical, and histological tests (**Figure 2b-f**). Control mice were treated with saline and euthanized after 1 day. To circumvent any unspecific inflammatory responses, residual endotoxin in the TmEnc formulation, a by-product of its recombinant production in *E. coli*, was reduced to safe levels (<0.03 EU/mL) before *in vivo* administration.

As expected, no gross outward changes in appearance (e.g., hunched posture, ruffled coat etc.) were observed in any of the mice post-injection, confirming that no acute inflammatory shock or severe hypersensitivity reactions had occurred as a result of TmEnc administration. There was also no significant weight loss in any of the TmEnc treated or control mice during the study; while normal weight gain was observed for mice in the 14-day group (**Figure 2b**).

To evaluate general health, serum samples obtained at each time-point were subjected to a comprehensive diagnostic panel that measured fourteen key electrolytes, enzymes, and proteins. When compared to controls, only modest changes to the serum levels of three known health markers were detected in TmEnc treated mice (see **Figure S3** for all acquired data sets). Following TmEnc administration, serum albumin (**Figure 2c**), a major blood component synthesized by the liver, was significantly reduced after 1 day (p=0.0404); alkaline phosphatase (ALP), an enzyme found in the liver and bone that breaks down proteins, and for which high levels can indicate liver disease/damage, was significantly lower 3 days later (p=0.0327) (**Figure 2d**); and alanine aminotransferase (ALT), an enzyme marker of liver injury, increased slightly after 1 day (**Figure 2e**). Importantly, all three of these analytes quickly recovered to relatively normal serum concentration ranges within the two-week study period, suggesting that, overall, TmEnc administration does not induce liver disease/damage.

In parallel, tissue sections of collected organs were subjected to hematoxylin and eosin (H&E) staining and examined in a blinded fashion (**Figure 2f** and **Figure S4**). No abnormal gross pathology, such as inflammation, necrosis, or hemorrhage, was detected in the liver, kidney, spleen, heart, or lung tissues of any treated animals, as compared to the saline controls, over the 14-day period. These results collectively suggest that the TmEnc nanocage does not adversely affect major organ function and integrity, making it a potentially safe NDDS candidate.

### Immune system interactions

When an NDDS enters the body, it interacts with the innate and adaptive immune systems, which not only affects its *in vivo* safety and tolerability, but also its biological fate and delivery efficiency.[38]

Toward this end, the immunostimulatory properties of TmEnc were first evaluated by assaying the obtained mouse serum samples for the circulating proinflammatory cytokines. Sera levels of tumor necrosis factor (TNF, <5 pg/mL) were slightly reduced in mice that received TmEnc (**Figure 3a**), however concentrations were below the detectable limit (<10 pg/ml). Similarly, both interleukin (IL)-6 and interferon gamma (IFNγ) concentrations were below the detectable limit (<10 pg/mL) in all treatment and control groups (**Figure 3b and c**). Both IL-6 and TNF levels here are below reported physiological levels. Thus, IV injected TmEnc does not appear to trigger any unsafe innate inflammatory responses.

**Figure 3.**
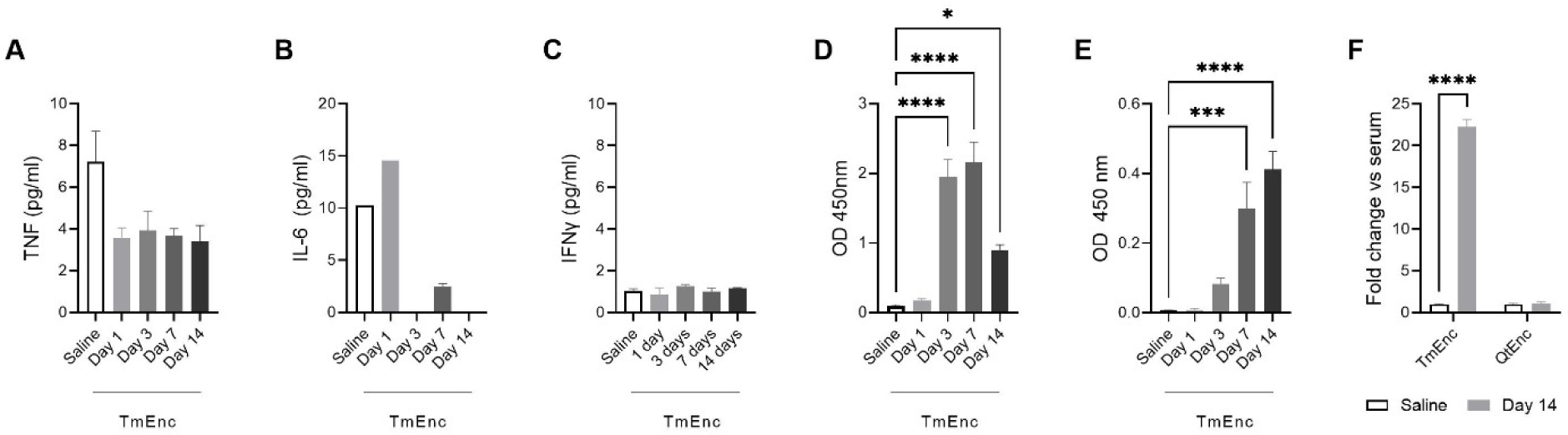
Immune responses to systemically administered TmEnc nanocages. Analysis of sera obtained from TmEnc treated mice determined the levels of circulating proinflammatory cytokines **(a)** TNF, **(b)** IL-6 and **(c)** IFNγ were below physiologically relevant concentrations. Results presented as mean ± SEM, one-way ANOVA, Tukey’s post hoc analysis, *n* = 4. Indirect ELISA detected the presence of anti-TmEnc **(d)** IgM and **(e)** IgG antibodies, confirming the immunogenicity of TmEnc. **(f)** Antibodies in the sera from TmEnc-treated mice showed no cross-reactivity with the QtEnc nanocage, demonstrating immune-orthogonality among encapsulin systems. Results presented as mean ± SEM, one-way ANOVA, Dunnett’s, *n* = 4 (*p≤0.05, ***p≤0.001, ****p≤0.0001).

We next investigated whether TmEnc induces any slower-acting adaptive immune responses, specifically antibody-mediated immunity (i.e., humoral immunity). Mice serum samples were therefore additionally analyzed for the presence of TmEnc-specific IgM (**Figure 3d**) and IgG (**Figure 3e**) antibodies by performing indirect ELISAs with plates coated with purified nanocages. As expected, sera from saline-treated control mice were negative for anti-TmEnc antibodies. In comparison, TmEnc-specific IgM antibodies were first detected in the sera of nanocage-treated mice 3 days after administration, before peaking at 7 days and then subsiding again by 14 days. In parallel, anti-TmEnc IgG antibodies were detectable after 3 days post-injection, and continued to increase significantly to a peak at 14 days. Together, these results show that TmEnc has immunogenic properties and triggers a classical adaptive immune response, wherein IgMs are initially produced by the body before isotype switching to high-affinity IgGs occurs, a process that takes approximately fourteen days to resolve. [39, 40] It is important to note that TmEnc-specific antibody generation implies the occurrence of an inflammatory response, which was most likely to have dissipated by the time pro-inflammatory cytokines were first measured in serum one day after TmEnc administration. Other IV injected PNCs have also been reported to induce the production of PNC-specific antibodies, and this immune response plays a key role in their elimination from the body.[41, 42] An elegant approach to lower the intrinsic immunogenicity of PNCs, is to identify B- and T-cell epitopes on their surfaces and alter and/or remove them through site-directed mutagenesis.[43] For example, the insertion of foreign peptides into the major immune region domains of human hepatitis B virus core protein (HBc) has been demonstrated to markedly reduce the PNC’s immunogenicity, aiding its subsequent application as a targeted NDDS.[44, 45]

The repeated administration of a PNC is known to further elevate antibodies levels, which makes them vulnerable to antibody-mediated neutralization and accelerated systemic clearance [42, 44–46]. Despite this knowledge, the generation of PNC-specific antibodies and the interactions they have with their target PNCs *in vivo* remains a relatively unexplored area. The Steinmetz lab highlighted the importance of understanding such phenomena by injecting the Potato virus X (PVX)-derived PNC into mice weekly, which led to increasing amounts of PVX-specific IgM and IgG antibodies.[46] Interestingly, intravital imaging revealed PVX-antibody complex formation and aggregation in the mouse vasculature, followed by isotype switching from IgM to IgG that resulted in reduced aggregate sizes. [47] Based on these unique observations, and our own finding that a single IV dose of TmEnc induces antibody production and IgM/IgG isotype switching over a two-week period, investigations into repeated administration of TmEnc is warranted.

To evade antibody-mediated neutralization and accelerated clearance upon repeated injection, researchers have proposed sequentially administering immune-orthogonal PNCs that have different protein sequences.[48] For instance, Ren and colleagues employed engineered PNCs derived from three different HBc capsids (i.e., human, woodchuck, and duck) that showed minimal antibody cross-reactivity between one another in mice initially immunized with wild-type human HBc [44]. Sequential administration of drug-loaded versions of these three PNCs resulted in lower immune clearance and more efficacious drug delivery in a pre-clinical cancer model.

Because encapsulins are prevalent throughout nature and also exhibit protein sequence diversity, we decided to explore their potential immune-orthogonality [49]. Here, the specificity of antibodies present in sera obtained from TmEnc-treated mice was assessed for cross-reactivity with the encapsulin derived from *Quasibacillus thermotolerans* (QtEnc). When compared to TmEnc (*T*=1, 60-mer, 24 nm), QtEnc (*T*=4, 240-mer, 42 nm) differs in size and structure, and shares only ∼20% amino acid sequence conservation (**Figure S5**).[50] Indirect ELISAs performed with plates coated with either TmEnc or QtEnc, confirmed both the presence of anti-TmEnc and, conversely, the complete absence of antibodies with specificity toward QtEnc (**Figure 3f**). This observed lack of antibody cross-reactivity between these two nanocages implies that immune-orthogonality exists among encapsulin systems, and could therefore be exploited to enhance their applicability in targeted drug delivery.

### Protein corona formation and characterization

When entering the bloodstream, a NDDS spontaneously adsorbs proteins, developing a surface layer referred to as the ‘protein corona’.[51] At first, weakly attached proteins form a ‘soft corona’ that is unstable and quickly exchange with proteins abundant in blood. Over time, proteins that bind more tightly to the NDDS establish a stable ‘hard corona’ coating.[51] The composition of the protein corona has a substantial impact on the size and surface properties of the NDDS, thus influencing how the body responds and processes it (e.g., immune interactions, biodistribution, and clearance).[52, 53]

We set out to characterize the hard protein corona formed on systemically administered TmEnc in order to better understand the encapsulin’s *in vivo* behavior. Given that we intended to visually track the *in vivo* biodistribution of TmEnc in downstream experiments (see section below), we elected to perform an *ex vivo* corona study with fluorescently-labelled encapsulin. Accordingly, the near-infra red (NIR) dye sulfo-Cy7-NHS was conjugated to lysine residues on the nanocage’s outer surface. In-gel fluorescence observed *via* SDS-PAGE confirmed the production of Cy7-labelled TmEnc (TmEnc^Cy7^) (**Figure 4a**), and TEM indicated no adverse changes to nanocage macrostructure (**Figure 4b**). UV–visible absorbance spectroscopy determined ∼50 dye molecules were conjugated to the surface of the TmEnc^Cy7^ nanocage (*data not shown*).

**Figure 4.**
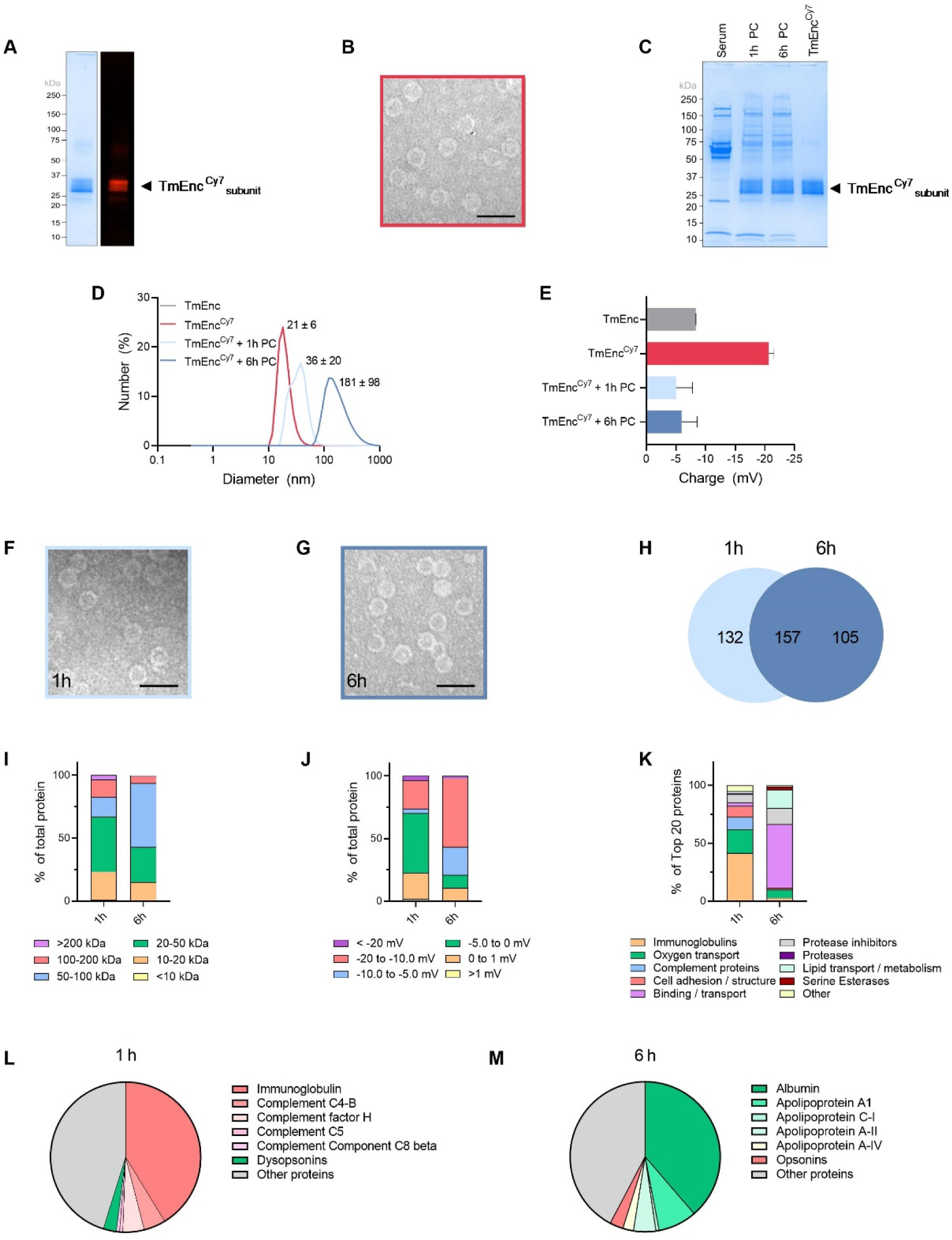
Protein corona formation on TmEnc^Cy7^ nanocages. **(a)** Coomassie staining and in-gel fluorescence observed SDS-PAGE confirmed Cy7 conjugation to TmEnc, resulting in fluorescent TmEnc^Cy7^.The TmEnc^Cy7^_subunit_ gel band appears thicker due to varying degrees of dye conjugation. **(b)** TEM verifying that dye-labelled TmEnc^Cy7^ retained its macrostructure (Scale bar = 50 nm). **(c)** SDS-PAGE visualizing the formation of hard protein coronas (PC) on TmEnc^Cy7^ after incubation with mouse serum for 1 h or 6 h. **(d)** DLS-measured size distributions of TmEnc^Cy7^ indicated size increases with protein corona coatings. **(e)** Zeta potential measurements revealed that the overall surface charge of TmEnc^Cy7^ increased from −21 mV to ∼5 mV upon protein corona formation; Results presented as the mean ± SEM, *n* = 3. TEM images showing the structure of TmEnc^Cy7^ was conserved following the formation of hard coronas after incubation with serum for **(f)** 1 h and **(g)** 6 h (Scale bars = 50 nm). **(h)** Venn diagram depicting the number of distinct serum proteins LC/MS identified in 1 h and 6 h coronas, and their respective overlap. Classification of the Top-20 most abundant proteins identified in the 1 h and 6 h coronas indicated differences in their composition, specifically: **(i)** molecular weight (kDa); **(j)** charge; **(k)** physiological function; and the **(l)** opsonin and **(m)** dysopsonin content.

Studies reporting corona formation on PNCs are sparse and tend to only focus on coronas that develop over short timeframes (≤ 1 hour). However, given that protein coronas on synthetic nanoparticles are known to change over time [51] it is important to understand how time might also affect corona formation and composition, as well as the subsequent fate of PNCs *in vivo* [54–56]. Taking this into account, TmEnc^Cy7^ was incubated with BALB/c mouse sera at 37°C for 1 h or 6 h. After washing away any weakly bound proteins, SDS-PAGE visually confirmed the presence of hard coronas on TmEnc^Cy7^ (**Figure 4c**).

DLS determined that the diameter of TmEnc^Cy7^ (21 ± 6 nm) was identical to unmodified TmEnc, but enlarged with corona development after 1 h (36 ± 20 nm), and even more so by 6 h (181 ± 98 nm) (**Figure 4d**). Zeta potential measurements revealed the overall surface charge of TmEnc^Cy7^ (−21 mV) was more negative than TmEnc (−8.4 mV) due to the neutralization of positive lysine residues following conjugation, but became more positive with the coronas (−5 mV) (**Figure 4e**). These changes in the size and charge of TmEnc^Cy7^ may not exclusively be attributed to corona coatings but also to some particle aggregation. TEM imaging further shows that corona decoration did not affect the structural morphology of the nanocage (**Figure 4f-g**). As depicted in **Figure 4h**, LC-MS analysis identified >250 different serum proteins within the coronas formed on the TmEnc^Cy7^, with 132 and 105 being unique to the 1 h and 6 h timepoints, respectively. This data shows that the surface of TmEnc^Cy7^ readily adsorbs serum proteins, forming hard coronas that alter their composition over time. This is intriguing as PNCs, unlike synthetic nanoparticles, typically exhibit minimal protein adsorption or complete protein avoidance in physiological fluids, resulting in low-density coronas or no corona formation at all.[56, 57]

The top-20 most abundant serum proteins adsorbed to TmEnc^Cy7^ after 1 h or 6 h (**Table 1**) were categorized according to their mass, charge, and physiological function (**Figure 4i-k**). Lower molecular weight proteins (<50 kDa) made up 67.0% of the 1 h corona, however, higher mass proteins (>50 KDa) became more prevalent over time, representing 57.0% of the 6 h corona (**Figure 4i**). At the pH of sera (pH 7.4), ≥70% the proteins within the coronas were negatively charged, with more strongly negative proteins (−20 to −10 mV) identified in the 6 h corona (55%) than the 1 h (22%) (**Figure 4j).** The classes of proteins adsorbed to TmEnc^Cy7^ differed markedly at each time point (**Figure 4k**). For instance, when compared to the original mouse serum, the 1 h corona was significantly enriched in immunoglobulins (IgGs; 41.3%) and complement proteins (10.7%), which are classed as opsonins; whereas the 6 h corona showed elevated levels of albumin (38.7%) and apolipoproteins (16.1%), which are considered dysopsonins.

**Table 1.**
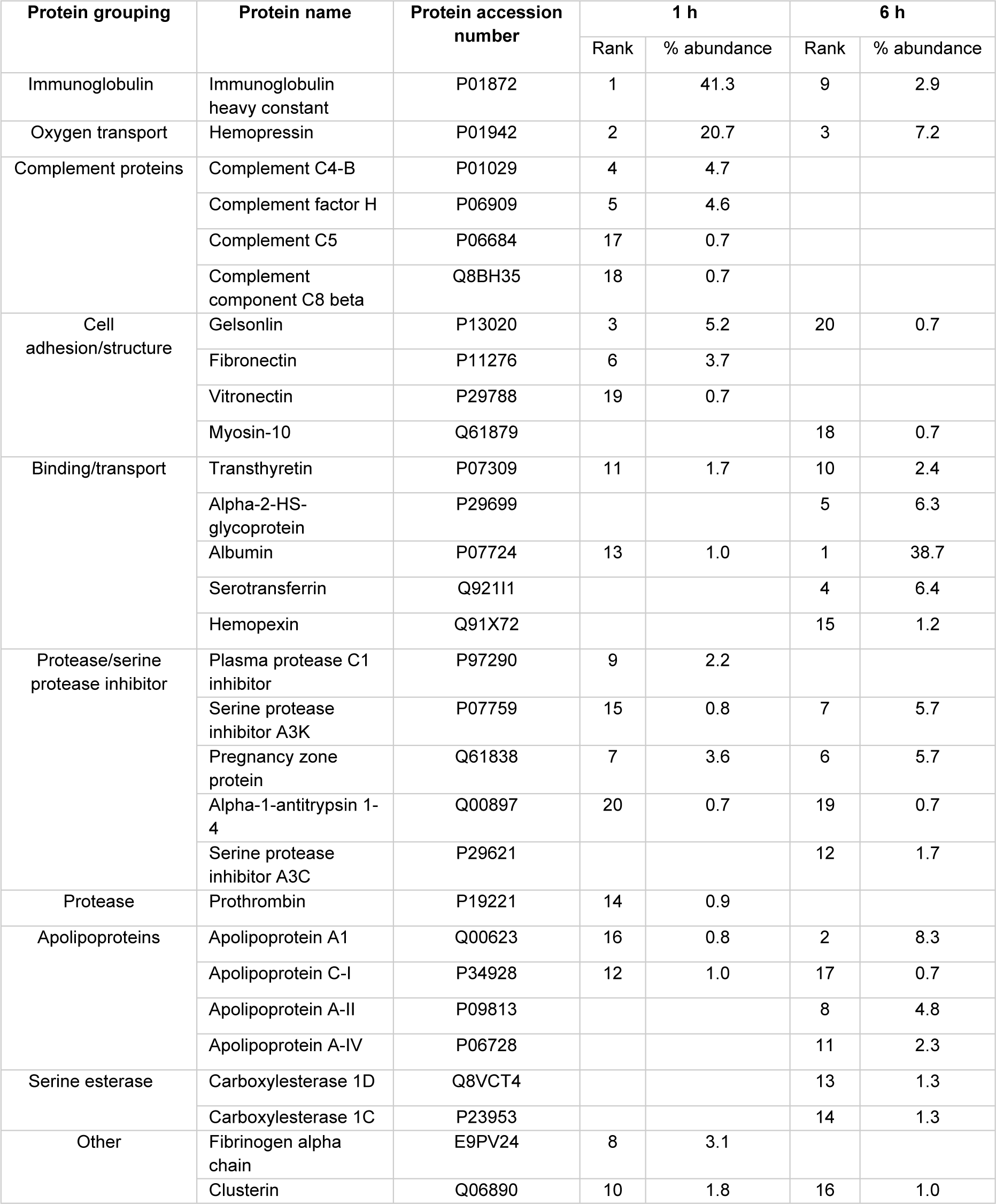
The Top-20 most abundant proteins identified in the TmEnc^Cy7^ hard coronas.

Upon deeper analysis, it was found that opsonizing proteins constituted 52.1% of the 1 h corona, but only 2.9% of the 6 h corona (**Figure 4l-m**). Opsonins participate in both the innate and adaptive immune responses.[58] When an NDDS acquires an opsonin-rich protein corona, it faces swift recognition and clearance by the mononuclear phagocyte system (MPS), an integral part of the innate immune system comprised of a network of phagocytic cells (e.g., monocytes and macrophages) found in the bloodstream, liver, and spleen.[59] The protein coronas of other PNCs have also exhibited a high abundance of opsonins, and their involvement in PNC clearance *via* the MPS has been implicated.[55, 56, 60–62]. In addition, opsonization can also initiate adaptive immune responses,[58] hence the rapid attachment of opsonins to TmEnc^Cy7^ supports our earlier observation that IV injected TmEnc induces the *in vivo* generation of nanocage-specific IgM/IgG antibodies (**Figure 3d-e**). To prevent such problematic opsonization and enhance their drug delivery efficacy, the outer surfaces of various PNCs have been coated with anti-fouling synthetic polymers (e.g., PEGylation)[36, 54, 63] or long repetitive hydrophilic peptides (e.g., PASylation, XTENylation) that reduce protein adsorption, and thus protein corona formation. [64, 65]

Unexpectedly, dysopsonizing proteins accounted for 54.7% of the 6 h corona, and only 2.8% of the 1 h corona (**Figure 4l-m**). This is an interesting observation because dysopsonins, unlike opsonins, typically hinder the phagocytosis of an NDDS by immune cells, allowing them to evade premature clearance from the body *via* the MPS.[58] Dysopsonizing proteins adsorbed on PNCs are relatively uncommon. However, the human ferritin PNC reportedly binds high levels of albumin when exposed to serum, which may help explain its relativity long blood circulation half-life in comparison to most other PNCs.[66] To help PNCs evade the immune system, researchers have pre-coated them with dysopsonizing albumin, or even modified them to selectively bind albumin during their systemic circulation in the body.[62, 67]

Similarly, because TmEnc appears to adsorb albumin onto its surface over time, the action of pre-coating it with albumin before administration could also prolong its blood circulation. While we acknowledge that capturing *in vivo* interactions within an *ex vivo* setting is challenging, our protein corona data does shed some light on TmEnc’s distinctive immune profile. The presence of an opsonin-rich corona after 1 h may have triggered an initial immune response, characterized by the production of pro-inflammatory cytokines and the initiation of antibody generation. However, a subsequent shift to a dysopsonin-rich corona over a 6 h period could have mitigated this acute inflammatory response, resulting in the reduced systemic levels of pro-inflammatory cytokines we observed in sera at 1-day post-injection (**Figure 3a-c**). Armed with our new insights into protein corona formation on TmEnc^Cy7^, we proceeded with evaluating the encapsulin’s biodistribution *in vivo*.

### *In vivo* biodistribution and clearance

In order to effectively deliver a therapeutic payload, an NDDS needs to localize and accumulate at a desired site-of-action (e.g., a solid tumor).[26, 29] With dye-labelled TmEnc^Cy7^ in hand (**Figure 4a-b**), we set out to visually track the general *in vivo* biodistribution of the nanocage (**Figure 5**). Herein, mice were IV injected with either TmEnc^Cy7^ or free Cy7 dye (control), mice groups (*n=3*) were then euthanized at 1, 3, 6, 12, 24, 48, or 72 h post-injection, and major organs subsequently harvested for *ex vivo* fluorescence imaging. Prior to administration, we confirmed that TmEnc^Cy7^ was highly stable when incubated with mouse sera *ex vivo*, with no unwanted protein degradation observed (**Figure S6**).

**Figure 5.**
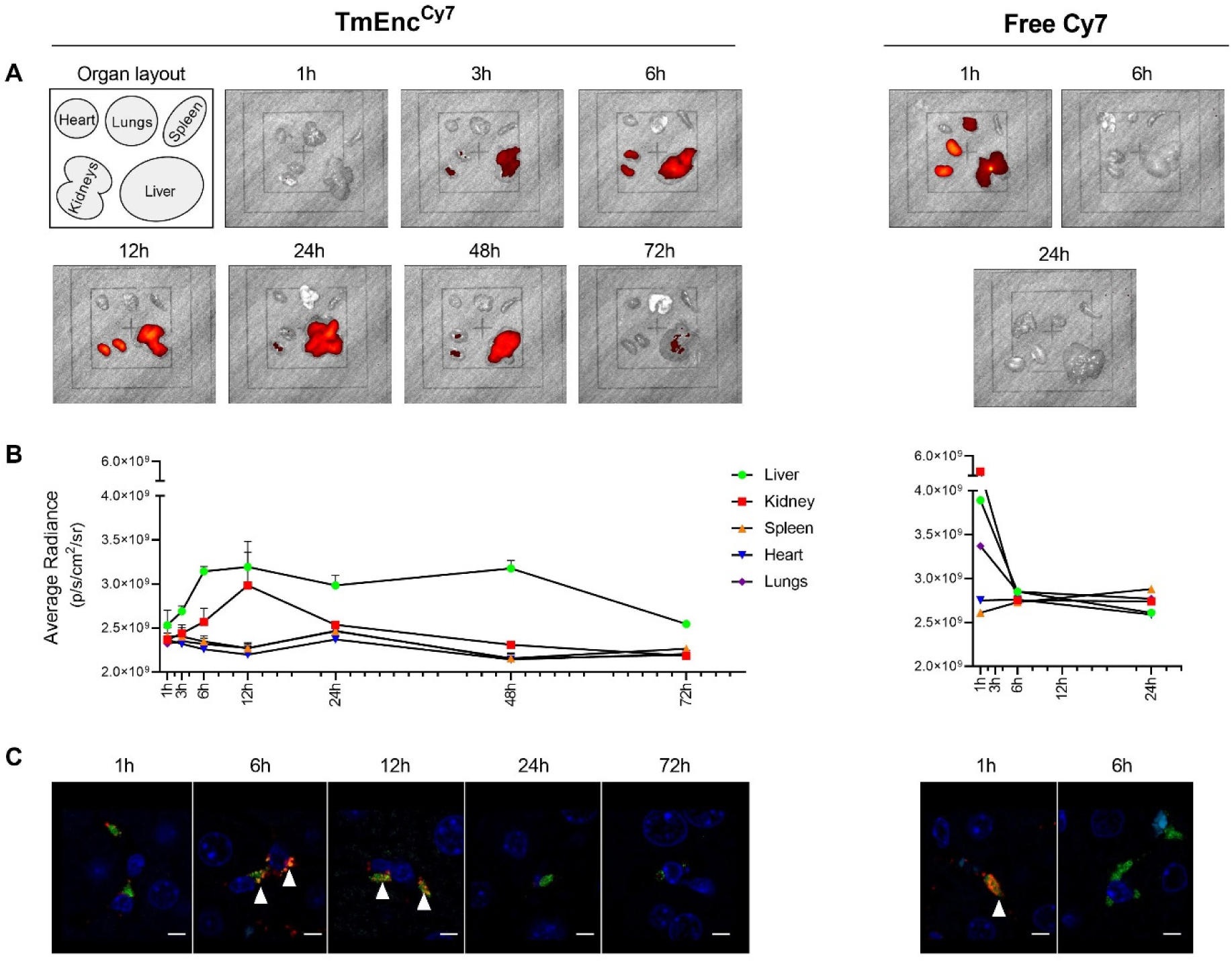
*In vivo* biodistribution and clearance of intravenously administered TmEnc^Cy7^. Fluorescent TmEnc^Cy7^ or free Cy7 dye was IV injected into BALB/c mice (2.5 mg/kg, *n* = 3), which were euthanized 1, 3, 6, 12, 24, 48 and 72 h after administration, with organs (heart, lungs, spleen, kidneys and liver) excised for fluorescence imaging. **(a)** NIR fluorescent images of organs showing the different biodistribution profiles of **(left panel)** TmEnc^Cy7^ and **(right panel)** free Cy7. **(b)** Quantitative analysis of NIR fluorescence intensity in the excised organs indicated that **(left panel)** TmEnc^Cy7^ predominantly accumulates in the liver by 6 h and then almost clears from the body within 72 h; **(right panel)** while free Cy7 is observed mostly in the kidneys within 1h but is rapidly eliminated by 6 h. Error bars represent SEM **(c)** Fluorescent confocal microscopy images of liver tissue sections showing that **(left panel)** TmEnc^Cy7^ was internalized by liver Kupffer cells within 6 h (white arrows), whereas **(right panel)** free Cy7 was up-taken by Kupffer cells after 1 h, but completely eliminated by 6h (For zoomed out microscopy images see **Supplementary Figure S5)**. Red = TmEnc^Cy7^; Green = CD68^+^ macrophage marker; Yellow = co-localization; Scale bars = 50 μm.

As presented in the NIR fluorescence images of excised organs (**Figure 5a**) and the corresponding quantitative analysis (**Figure 5b**), TmEnc^Cy7^ accumulated primarily inside the liver, followed by the kidneys, and only negligible amounts were found in the spleen, lungs, and heart. Specifically, mice that received TmEnc^Cy7^ began to display liver deposition 3 h after injection, which steadily increased to a peak at 6 h, followed by gradual clearance by 72 h. Concurrent TmEnc^Cy7^ accumulation in the kidneys was also detected 3 h after injection, reaching an eventual peak at 12 h, and then quickly clearing by 24 h. In contrast, free Cy7 dye mostly accumulated within the kidneys 1 h post-injection, with some moderate deposition observed within the liver and lungs. By 6 h, free Cy7 was rapidly cleared through the renal system, likely due to the dye’s small molecular size.[68] The near-complete deposition of TmEnc^Cy7^ within the liver over a 6 h period is not unexpected, as IV administered PNCs, like most NDDSs, ordinarily accumulate in MPS organs (i.e., liver and spleen).[36, 54, 69] Indeed, other PNCs have shown much faster MPS organ deposition than TmEnc^Cy7^; for instance, over 90% of injected Cowpea mosaic virus capsid has been detected inside the liver in under 30 mins.[69]

The biodistribution profile for TmEnc^Cy7^ implies that the encapsulin is sequestered from the blood circulation by the liver MPS. To probe these interactions between TmEnc^Cy7^ and the liver MPS, we prepared immunostained liver tissue sections for Kupffer cells (CD68^+^), resident liver macrophages and the principal cells of the MPS. As depicted in **Figure 5c**, fluorescence confocal microscopy revealed the entry of TmEnc^Cy7^ into the liver sinusoids within 1 h post-administration (**Figure S7**). By 6 h, the nanocages were internalized by Kupffer cells, and to a lesser extent, the hepatocytes. TmEnc^Cy7^ internalized by Kupffer cells began to degrade after 24 h, with only negligible amounts present at 72 h. No nanocages were visualized inside hepatocytes by 24 h, which implies partial clearance *via* hepatocytes and the hepatobiliary system. On the other hand, some free Cy7 was observed in Kupffer cells after 1 h, but was completely eliminated by 6 h. Although PNC sequestration by the liver MPS is well-documented, only a handful of studies have directly visualized the internalization of IV administered PNCs by Kupffer cells.[36, 70, 71] Similar to our own findings, all of these reports see Kupffer cell uptake within 1-4 hours of injection.

Based on these results, TmEnc^Cy7^ is predominantly sequestered from systemic circulation by the liver MPS within 6 h, after which it is taken up by Kupffer cells and gradually degraded over a period of 72 h. This is consistent with our finding that TmEnc^Cy7^ rapidly acquires a serum-derived protein corona rich in opsonins capable of marking an NDDS for MPS recognition and clearance. Importantly, the preceding safety profile of encapsulin indicates that the observed deposition of the nanocage within the liver is not associated with any hepatic toxicity.

## 3. Conclusions

In this pre-clinical work, we assessed the interactions between an IV-administered TmEnc and the key biological barriers encountered by NDDSs, and their influence on the encapsulin’s *in vivo* fate. Our purified TmEnc formulation was monodisperse, colloidally stable, storable, and non-hemolytic. In mice, IV injected TmEnc exhibited an excellent safety profile, with no adverse changes in animal weight, nor the serum levels of toxicity biomarkers, although there were some transient changes in liver function markers. Moreover, there were no signs of gross pathological changes in any major organs. On the other hand, our results revealed that the encapsulin has immunogenic properties, and interacts with both the innate and adaptive arms of the immune system. Here, TmEnc-specific IgM and IgG antibodies were detected in mouse sera, indicating an antibody-mediated immunity response to the nanocage. This also correlated with the nanocage quickly developing a protein corona with a high abundance of opsonins, which enables the PNC to be recognized by the immune system. Finally, the TmEnc’s *in vivo* biodistribution profile showed that it was removed from blood circulation by the MPS of the liver, which culminated in its uptake and biodegradation by Kupffer cells.

The apparent interactions between TmEnc and the immune system has implications for the development of encapsulins into targeted NDDSs. Premature immune clearance will likely prevent meaningful amounts of an encapsulin-based delivery system reaching an intended site-of-action, limiting its therapeutic effect. Similarly, the antibody-mediated immune response to TmEnc suggests that encapsulins may also be susceptible to accelerated systemic clearance upon repeat administration. Further detailed research is underway in our lab to ‘stealth’ encapsulins to evade immune recognition and clearance. We envisage that many surface coatings employed to stealth other PNCs will be compatible with encapsulins e.g., synthetic polymers, [36, 54, 63] biological polymers,[64, 65] and self-proteins [62]. Other alternative and/or complementary stealth-ing strategies could involve site-directed removal of specific immunogenic regions (i.e., epitopes), as well as exploiting the apparent immune-orthogonality that was observed between different encapsulin systems in this study.[44, 45, 49]

Despite being a relatively new class of PNC, encapsulins have already been re-engineered to possess properties and functions that directly and indirectly lend themselves to drug delivery. These include alternative mechanisms for packaging diverse cargoes (e.g., proteins, nucleic acids, synthetic molecules);[12, 72, 73] integration of surface coupling systems (e.g., spytag/spycatcher, split-inteins) for the modular display of cell-targeting ligands;[10, 15, 17] as well as altered subunit interfaces that permit controlled payload release.[74] Ultimately, we anticipate that our valuable insights into TmEnc’s *in vivo* behavior and fate, combined with ongoing advances in encapsulin engineering, will inform and expedite the translation of these unique PNCs into safe and effective drug delivery systems.

## 4. Experimental Section

### Materials

All chemicals and reagents used in this study were purchased from Sigma-Aldrich or ThermoFisher Scientific, unless stated otherwise.

### Protein nanocage production and characterization

*TmEnc nanocage production and purification*: *E. coli* BL21 (DE3) cells (New England Biolabs) harbouring a pETDuet-1 expression plasmid containing the codon-optimised TmEnc gene (UniProt: TM_0785) was cultured in Luria-Bertani (LB) medium supplemented with carbenicillin (100 μg/mL).[5] Starter cultures were grown overnight at 37°C and used to inoculate flasks of 500 mL LB media (1:100 v/v). Cultures were grown aerobically at 37°C until 0.5-0.6 OD_600_ was reached. Protein expression was then induced by the addition of 0.1 mM isopropyl-β-d-thiogalactopyranoside (IPTG). Cells were harvested by centrifugation (9,000 *g*, 10 min, 4°C) and the pellets stored at −30°C until further use.

The pellet from 1 L of cell culture was resuspended in Tris buffer (5 mL/g wet cell mass) (20 mM Tris, 150 mM sodium chloride (NaCl), pH 7.5) containing lysis components (1.5 mM MgCl_2_, 25 U/mL Benzonase nuclease, Roche Complete™ Mini, EDTA-free Protease Inhibitor Cocktail (one tablet per 30 mL)). Cell lysis was performed by probe sonication on ice at 50% amplitude for 10 sec on, 20 sec off, for 5 min, repeated 4 times. Cellular debris was then removed by centrifugation at 10000 *g*, 4°C for 15 min. The supernatant was next decanted and incubated on ice for 30 min to enhance Benzonase nuclease digestion. To denature host proteins and partially purify the recombinant nanocages, the soluble protein fraction was heat-treated at 65°C for 20 min and subsequently centrifuged at 10000 *g*, 4°C for 15 min. Soluble encapsulins in the resulting supernatant were precipitated by the addition of polyethylene glycol 8000 (PEG 8000) (8% w/v) and NaCl (0.5 M final concentration) followed by incubation on ice for 30 min. Precipitated proteins were then centrifuged at 10000 *g*, 4°C for 15 min, with the resulting protein pellet resuspended in Tris buffer and sterilized using 0.45 μm and 0.2 μm syringe filters (Merck, US).

The resulting protein solution was subjected to size exclusion chromatography (SEC) using a HiPrep™ 26/60 Sephacryl S-500 HR column (Cytiva, USA) at a flow rate of 1 mL/min. Fractions containing assembled encapsulin were determined by SDS-PAGE and Native-PAGE then pooled and loaded onto anion-exchange chromatography (AEX) column HiPrep Q FF 16/10 (Cytiva, USA) at a flow rate of 3 mL/min. Proteins were eluted by stepwise gradient of 0-1M NaCl in Tris buffer. Fractions containing encapsulin were combined and concentrated by Vivaspin 20 (Sartorius) spin filters (100 KDa cutoff) and frozen at −20°C until further use. Purified encapsulin concentration was determined by a Bradford assay.

#### Polyacrylamide Gel Electrophoresis (PAGE)

Purified protein samples were visualized to evaluate purity by SDS-PAGE using the Bio-Rad mini-protean system (Bio-Rad laboratories). Samples were mixed in a 1:1 ratio with 2x Laemmli sample buffer containing 50 mM 1,4-dithiothreitol (DTT) and heated at 99°C for 10 min before loading onto the gel. This was then run for 35 min at 200 V on a 4-20% polyacrylamide gel (MiniPROTEAN TGX, BioRad) in 1x SDS running buffer (25 mM Tris, 192 mM glycine, 1% (w/v) SDS, pH 8.3). Gels were imaged using the Cy7 setting (Excitation: Red Epi illumination, Emission: 700/50 filter) for fluorescence on Bio-Rad Chemidoc MP imager and subsequently stained for proteins using Coomassie R-250.[75]

#### Dynamic Light Scattering (DLS)

The zeta-potential, PDI, and hydrodynamic diameter of purified encapsulins were measured on the Malvern Zetasizer Nano ZS. Three measurements were performed at 25°C in ZEN040 cuvettes or capillary cells containing 70 µL of sample diluted in PBS (pH 7.4) to a final concentration of 0.15 mg/mL. Data analysis was performed in Zetasizer Nano software.

#### Transmission Electron Microscopy (TEM)

For the visualization of encapsulin self-assembly and protein corona formation, TEM was performed using a Philips CM10 microscope operating at 100 kV. 10 µL of sample (100 µg/mL) was deposited onto carbon film coated 300 mesh copper grids (ProSciTech) and negatively stained with uranyl acetate replacement stain (UAR-EMS) (1:3 dilution) for 1 h, washed with ultrapure water and allowed to dry for at least 24 h.

#### Endotoxin Removal

To remove endotoxin from purified TmEnc prior to *in vivo* studies, a Triton X-114 phase separation method was employed.[76] Initially, 1% (v/v) Triton X-114 was added to the sample and incubated with agitation for 15 min at 4°C. Subsequently, the samples were incubated at 37°C for 5 min and centrifuged at 10000 *g* for 5 min at 37°C to separate the two phases. The nanocage-containing supernatant was carefully collected and adjusted back to the original volume. This phase separation process was repeated twice. To eliminate any residual Triton X-114, Bio-beads SM-2 Resin (Bio-Rad) were introduced to the purified nanocage samples at a ratio of 5 g per 25 mL. The mixture was incubated with agitation for 2 h at room temperature. After incubation, the samples were centrifuged at 5000 *g*, 5 min at room temperature, and the resulting supernatant collected. To verify the removal of endotoxin from the encapsulin formulations, the collected supernatant was subjected to a Limulus Amebocyte Lysate gel clot test (Fujifilm) with a sensitivity of 0.03 EU/mL.

#### Dye labelling

Sulfo-Cyanine 7-NHS ester (sulfo-Cy7) was purchased from Lumiprobe. For amine-NHS coupling, TmEnc solutions (∼2 mg/mL) were prepared in 100 mM sodium bicarbonate buffer (pH 9.0). Sulfo-Cy7 was mixed with TmEnc at a molar ratio of 8:1, and mixed *via* gentle agitation for 24 h at room temperature. Following the reaction, excess dye was removed *via* benchtop SEC using sephadex G50 (Sigma), eluted with 1x PBS (pH 7.4). Fractions containing dye conjugated encapsulin were pooled and concentrated. To calculate the degree of labelling (DOL), the molar protein concentration was first calculated using the protein absorbance at 280 nm and 750 nm (A_max_), correction factor (0.04) and ε of protein. Moles of dye per mole protein was then calculated using the molar protein concentration, A_max_ and ε of the dye.

#### Protein Stability

To test the storage stability of TmEnc, 100 µL aliquots of encapsulin (10 µM in 20 mM Tris-HCl, pH 7.5) were subject to 1 or 4 freeze-thaw cycles, followed by TEM, DLS, and SDS-PAGE densitometry analysis to assess their stability. Freeze-thaw cycles involved storing the sample at −80°C for 20 min, followed by thawing in water at room temperature. Samples were centrifuged at 17,000 *g* for 30 min at 4°C to remove any aggregates before analysis. Sample that had not been frozen was defined as 100% soluble.

To assess the stability of TmEnc^Cy7^ in mouse serum, samples were diluted to 100 ug/mL in 1x PBS (pH 7.4) and mixed with 55% mouse serum. Samples were then vortexed briefly before incubation at different time points over 24 h (0, 1, 3, 12 and 24 h). SDS-PAGE analysis was performed as described above.

#### Hemolysis Assay

TmEnc was tested for toxicity *via* a haemolysis assay. In this assay, the amount of hemoglobin released by red blood cells (RCBs) following co-incubation of RBCs and encapsulins was determined. Briefly, whole blood was collected, the plasma layer removed, and RBCs washed with PBS. RBCs were resuspended in PBS (pH 7.4), then diluted 1:50 in PBS for the assay. 10 µl of TmEnc was added with 190 µl of diluted RBC solution added. 10 µl of 20% Triton X-100 or PBS were used as the positive and negative controls, respectively. The plate was incubated at 37°C for 1 h, then centrifuged at 500 *g* to pellet intact RBCs. 100 µl of supernatant was removed to a new plate, and absorbance was read at 400 nm. Results are presented as percentage hemolysis as compared to the Triton X-100 control.

#### Protein corona preparation and analysis

To evaluate protein corona formation, 112.5 µL of pure TmEnc in 50 mM HEPES (pH 7.4) was added to 55% of mouse plasma (Final volume of 250 µL) and incubated at 37°C (in a water bath) for 1 h and 6 h. Unbound plasma proteins were separated from TmEnc nanoparticles by sucrose cushion ultracentrifugation. Briefly, after incubation, the samples were placed onto a 6 mL sucrose cushion (25% w/v sucrose, 50 mM HEPES, pH 7.4) in 10.4 mL polybottle ultracentrifuge tubes and HEPES buffer was added to completely fill the tubes. The samples were centrifuged for 2 h at 160,000 *g* at 4°C. Carefully, without taking the pellet, the supernatant was then removed, and the pellet was resuspended in 250 µl HEPES buffer and sucrose gradient was performed once more to wash unbound proteins. The washed pellet was then collected (resuspended in 250 µl HEPES buffer) and stored at −20°C until required for mass spectrometry preparation.

#### Liquid chromatography mass spectrometry (LC-MS)

For mass spectrometry analysis, the protein content of the recovered corona-encapsulin complexes was quantified using a Bradford assay following the manufacturer’s instructions. Normalized samples were incubated with the same volume of 10% Sodium deoxycholate (SDC) and 1:50 to final concentration of 5 mM (tris(2-carboxyethyl) phosphine) (TCEP) and 2-iodoacetamide (IAA) and incubated for 10 mins at 95°C. Afterward, samples were incubated overnight at 37°C with sequencing grade modified trypsin (0.01 mg/mL; Promega Corporation) to allow protein digestion. The digestion reaction was stopped by adding 10x volume of 90% acetonitrile (ACN) and 1% Trifluoroacetic Acid (TFA) and centrifuged to pellet and insoluble proteins. The digested peptides were loaded onto solid phase extraction (SPE) columns. Columns were first equilibrated by adding 90% ACN and 1% TFA, then samples were added and centrifuged at 4,900 *g* for 2 min, until all the samples passed through the SPE. Samples were washed twice with 100 μl of 10% ACN and 0.1% TFA then eluted by adding 50 μL of elution buffer (71 μL 1M NH_4_OH_3_, 800 μL of 100% ACN, 129 μL Water). The supernatant containing the peptides was collected and dried using a speed vacuum for about 2 h. Samples were then resuspended in 25 μL of 2% ACN 0.2% TFA in water, and 1 μL samples were loaded into a Q Exactive Plus hybrid quadrupole− orbitrap mass spectrometer (ThermoFisher) using Acclaim PepMap 100 C18 LC columns (ThermoFisher). Samples were analyzed using the software PEAKS Studio 8.5 (Bioinformatics Solutions Inc.), and proteins were identified using the manually reviewed UniProtKB/SwissProt database. Only proteins identified by at least one unique peptide were included in the analysis.

### Animal experiments

#### Ethical statement

All animal models used were approved by The University of Technology Sydney’s Animal Care and Ethics Committee (ETH22-6953) and were in accordance with the Australian Code for the Care and Use of Animals for Scientific Purposes, 8th Edition, 2013 guidelines.

#### Animal Husbandry

Male BALB/c mice 8 weeks old (20-26 g) were kept on a 12 h light/dark cycle with food and water available *ad libitum*. To assess toxicity of TmEnc, mice received one intravenous injection *via* the tail vein of either saline or TmEnc (5 mg/kg). Mouse weight was tracked daily. Mice were euthanized at 1, 3, 7, and 14 days post-injection *via* cardiac puncture under anesthesia. Tissues collected were halved, and either snap frozen in liquid nitrogen, or fixed in paraformaldehyde. Whole blood was collected and allowed to clot for 15 min at room temperature, then centrifuged (1,500 *g*, 5 min at 4°C), and the serum layer removed.

#### Biodistribution

To determine the biodistribution of the TmEnc to the tissues, dye TmEnc^Cy7^ (2.5 mg/mL) was injected intravenously. Mice were euthanized at 1, 3, 6, 12, 24, 48, and 72 h post-injection. Following euthanasia, organs were excised and tissue accumulation was visualized using the IVIS imaging system (Caliper Life Sciences).

#### Serum Analysis

Serum samples were assessed for albumin, alkaline phosphatase, alanine aminotransferase, amylase, total bilirubin, blood urea nitrogen, calcium, phosphate, creatine, glucose, sodium, potassium, total protein and globulin using a VetScan VS2 Chemistry Analyzer (Zoetis Inc). Circulating levels of IL-6 and TNF were determined using a cytokine bead array (BDTM Cytometric Bead Array Mouse IL-6/TNF Flex Set, BD Biosciences). Samples were analyzed in technical triplicates.

#### ELISA

To determine amounts of TmEnc-specific IgG or IgM in mouse sera, 96-well Nunc MaxiSorp Immuno Plates (Thermo Fisher Scientific) were coated with 2.5 µg/ml TmEnc nanocages, or QtEnc for immune orthogonality studies, (100 µl per well) in PBS overnight at 4°C. Unbound TmEnc was removed from the wells, which were then washed 3 times with 200 µl PBST buffer (PBS + 0.05% Tween 20). Wells were then blocked with protein block (3% BSA fraction V (Merck) or 10% FBS) in PBS for 1 h at room temperature, followed by washing as above. Next, sera from TmEnc treated mice was diluted in PBS (1:1000 for IgG, 1:200 for IgM) then added to the wells and incubated for 1 h at room temperature. Wells were washed as above, then 50 µl of TMB Chromogen Solution (Thermofisher) was added. The reaction was stopped with the addition of 50 µl of 6% H_2_PO_4._ Absorbance was finally measured at 450 nm using a Tecan plate reader.

#### Histology

Following excision, tissues were fixed in 4% paraformaldehyde, processed, and embedded in paraffin wax. Paraffin embedded organs were sectioned at 5 µm. Slides were rehydrated and stained with H&E, dehydrated, mounted and cover slipped. To assess pathology, stained slides were scanned using a Zeiss AxioScan slide scanner. Each scanned image was examined by two independent scorers using a semi-qualitative scoring system to identify inflammatory, fibrotic, vascular or necrotic changes (**Figure S4**), where: 0 = no abnormality seen; 1 = mild; 2 = moderate; and 3 = severe histological abnormality seen. Specific abnormalities were recorded for each section.

To visualize the internalization of TmEnc^Cy7^ by Kupffer cells, liver tissue sections were stained for the macrophage marker CD68. Briefly, rehydrated slides underwent antigen retrieval in citrate buffer, were blocked in phosphate buffer with goat serum and BSA at room temperature for 15 min, then incubated in anti-CD68 (1:500, Abcam) for 2 h at room temperature. Slides were then washed three times for 3 min in PBST and incubated in an anti-rabbit AF488 (1:2000, Invitrogen) and DAPI (1:500) cocktail for 2 h at room temperature. Slides were then washed as pervious, mounted and cover slipped.

#### Statistical analysis

Statistical significance was determined *via* Ordinary One-way ANOVA with Dunnett’s or Tukey’s post-hoc analysis using GraphPad PRISM. Statistical significance of ranked data was determined *via* Kruskal-Wallis test using GraphPad PRISM. Results are presented as mean ± standard error of mean (SEM).

## ASSOCIATED CONTENT

### Supporting Information

Chromatograms of TmEnc nanocage purification; SDS-PAGE of TmEnc’s storage stability; serum analyte measurements for general health assessment; pathology scoring of tissues stained with H&E; protein sequence and structure alignments of TmEnc and QtEnc, SDS-PAGE of TmEnc^Cy7^’s stability in serum; confocal fluorescent microscopy images visualizing TmEnc^Cy7^ uptake by liver cells.

## AUTHOR INFORMATION

### Author Contributions

C.R. co-designed research, conducted animal studies and *in vitro* bioassays, performed data analysis, wrote manuscript. C.S. protein production and purification, protein bioconjugation, protein corona isolation, wrote manuscript. I.B. protein stability, TEM imaging, revised manuscript. D.D. protein corona preparation, revised manuscript. C.A.G. performed histological analysis, revised manuscript O.V. co-designed study, revised manuscript. L.C.P. co-designed study, revised manuscript. A.C. conceptualized and co-designed research, assisted animal studies, supervised project, and wrote manuscript.

### Notes

The authors declare no competing financial interest.

## ACKNOWLEDGMENTS

This research was supported by a Priority-driven Collaborative Cancer Research Scheme grant (1182082) co-funded by Cancer Australia, The Kids’ Cancer Project, and the Australian Lions Childhood Cancer Research Foundation. This work was additionally supported by grants from Dementia Australia Research Foundation, Mason Foundation, and the National Foundation for Medical Research and Innovation. A.C. is supported by a Chancellor’s Research Fellowship from the University of Technology Sydney (UTS). I.B. is supported by a Dementia Australia Research Foundation PhD scholarship. The authors acknowledge the technical and scientific assistance of UTS’ Proteomics, Lipidomics and Metabolomics Core Facility, and also the Ernst Animal Facility. The authors also pay their respects to the Gadigal people, who are the traditional custodians of the land on which this research took place.

