## Supplementary information for "*In vivo* fate of systemically administered encapsulin protein nanocages and implications for their use in targeted drug delivery"

Contents

**Supplementary Figure S1.** Chromatograms of TmEnc nanocage purification

**Supplementary Figure S2.** Storage stability of the TmEnc nanocage

**Supplementary Figure S3.** Serum analytes investigated for general health assessment

**Supplementary Figure S4.** Pathology scoring of tissues stained with H&E

**Supplementary Figure S5.** The protein sequence and structure of *T*=1 TmEnc differs from *T*=4 QtEnc

**Supplementary Figure S6.** Stability of dye-labelled TmEnc^Cy7^ in mouse serum

**Supplementary Figure S7.** Visualisation of TmEnc^Cy7^ uptake by liver cells

**References**

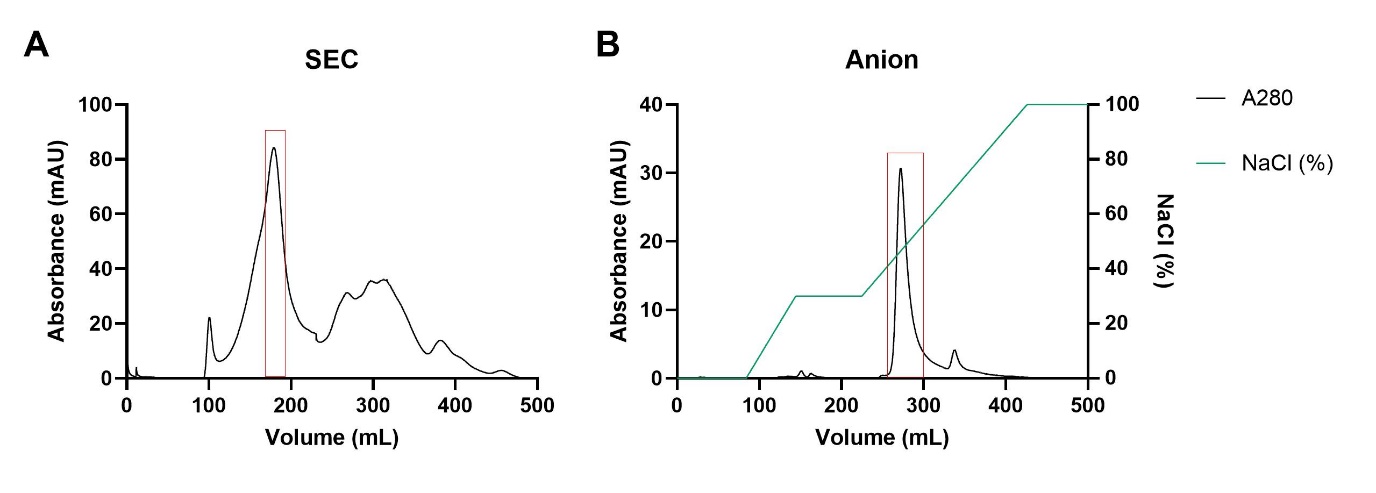

**Figure S1. C****hromatograms of TmEnc nanocage purification. (a)** Size exclusion chromatography (SEC) of TmEnc. The protein of interest (red rectangle) eluted between 170-190 mL with absorbance at 280nm (black line). **(b)** Anion exchange chromatography of SEC-purified TmEnc eluted at 40-50% NaCl (blue line).

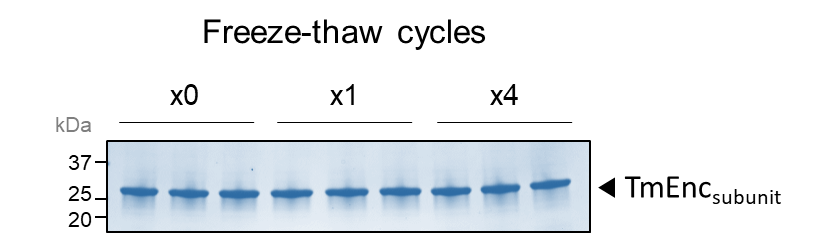

**Figure S2.** **Storage stability of the TmEnc nanocage.** Purified TmEnc underwent 0x, 1x, or 4x freeze-thaw cycles at -80°C. SDS-PAGE analysis shows that the recovered soluble TmEnc protein was >99% after each freeze-thaw.

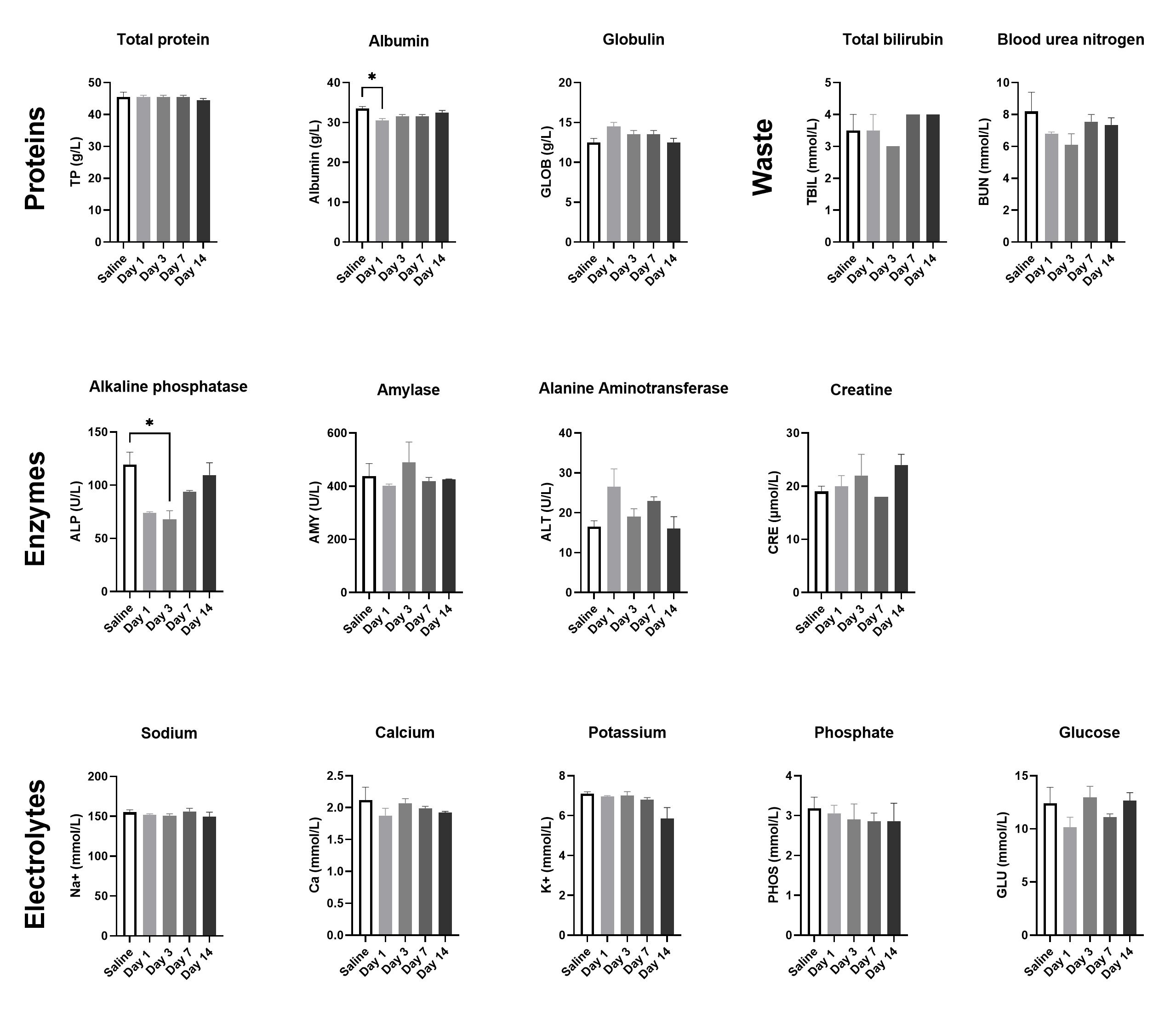

**Figure S3. Serum analytes investigated for general health assessment.** 5 mg/kg of TmEnc was administered to BALB/c mice *via* tail-vein injection*.* At 1, 3, 7, or 14 days post-administration, sera was collected and assayed for markers of general health using the VetScan VS2 Chemistry Analyser (Zoetis Inc). Results are presented as mean ± SEM. Statistical significance was determined via one-way ANOVA with Tukey’s post-hoc analysis, *n*=2 (2x pooled samples from 2x animals), *P<0. 05.

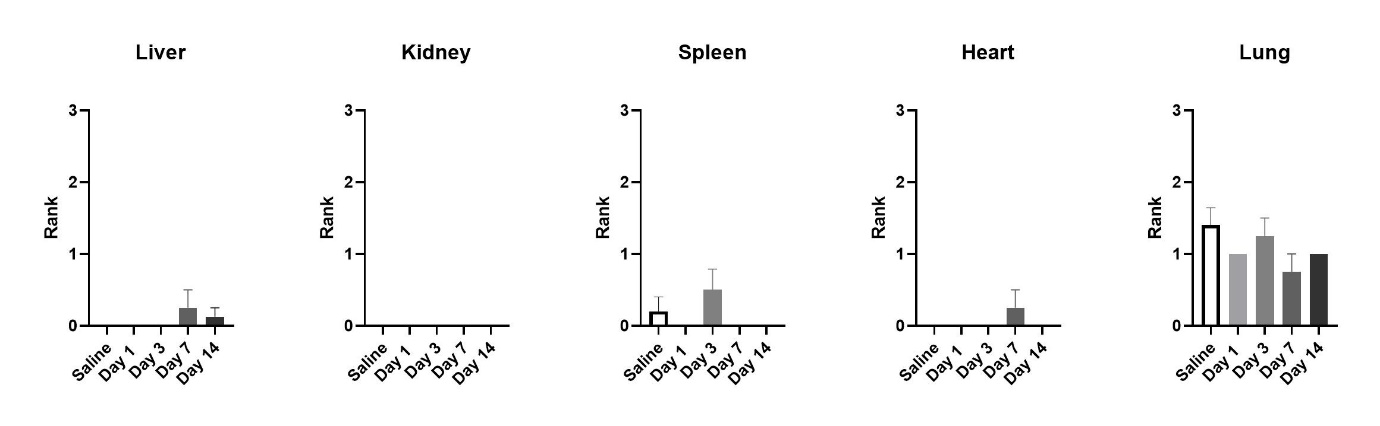

**Figure S4. Pathology scoring of tissues.** Sections were stained with H&E, scanned using an AXIO Slide scanner and were scored for pathology. Results presented as mean ± SEM. Statistical significance was determined *via* Kruskal-Wallis test. *n* = 4.

**H&E scoring:** Each scanned image was examined by two independent scorers using a semi-qualitative scoring system to identify inflammatory, fibrotic, vascular or necrotic changes where: 0 = no abnormality seen; 1 = mild; 2 = moderate; and 3 = severe histological abnormality seen. Specific abnormalities are recorded for each section.

Liver morphology appeared to be consistent for species and predominantly normal. Several sections, regardless of saline or TmEnc administration, exhibited discrete patches of binucleated cells with eosinophic cytoplasm which may represent normal age-related changes in these mice. One section had a small area of pale cells that may represent necrosis, but this was not extensive. Occasional small areas with extravascular erythrocytes were noted, but there were no areas of overt or extensive haemorrhage.

Kidney morphology appeared to be consistent for species and predominantly normal. Occasional small areas with extra vascular erythrocytes were noted, but there were no areas of overt or extensive haemorrhage.

Heart morphology appeared to be consistent for species and predominantly normal. Occasional small areas with extravascular erythrocytes were noted but there were no areas of overt or extensive haemorrhage. One section had a small cluster of lymphocytes and a small cluster of large pale cells with prominent nuclei in the surrounding adipose tissue.

Lungs generally appeared to contain large numbers of erythrocytes and in 2 sections, one mouse each from saline and TmEnc Day-3, this was quite extensive. The alveoli spaces were quite congested/collapsed in the majority of sections, but 5 sections showed a normal appearance of alveoli. In line with this, similar changes were evident in saline-treated animals, and is therefore, unlikely due to encapsulin administration. This appearance is most likely due to the lack of perfusion and inflation following dissection. Lungs were removed following cardiac bleed and immersion fixed in 4% paraformaldehyde. Oesophagus, skeletal muscle, inflammatory cells (BALT) and white and brown adipocytes were noted in the surrounding tissue in 3 sections. Epithelium of airways and blood vessels appeared normal.

Spleen morphology appeared to be mostly consistent for species with distinct white and red pulp and was predominantly normal. There were occasional megakaryocytes seen in the red pulp.

Spleen morphology appeared to be mostly consistent for species with distinct white and red pulp and was predominantly normal. There were occasional megakaryocytes seen in the red pulp.

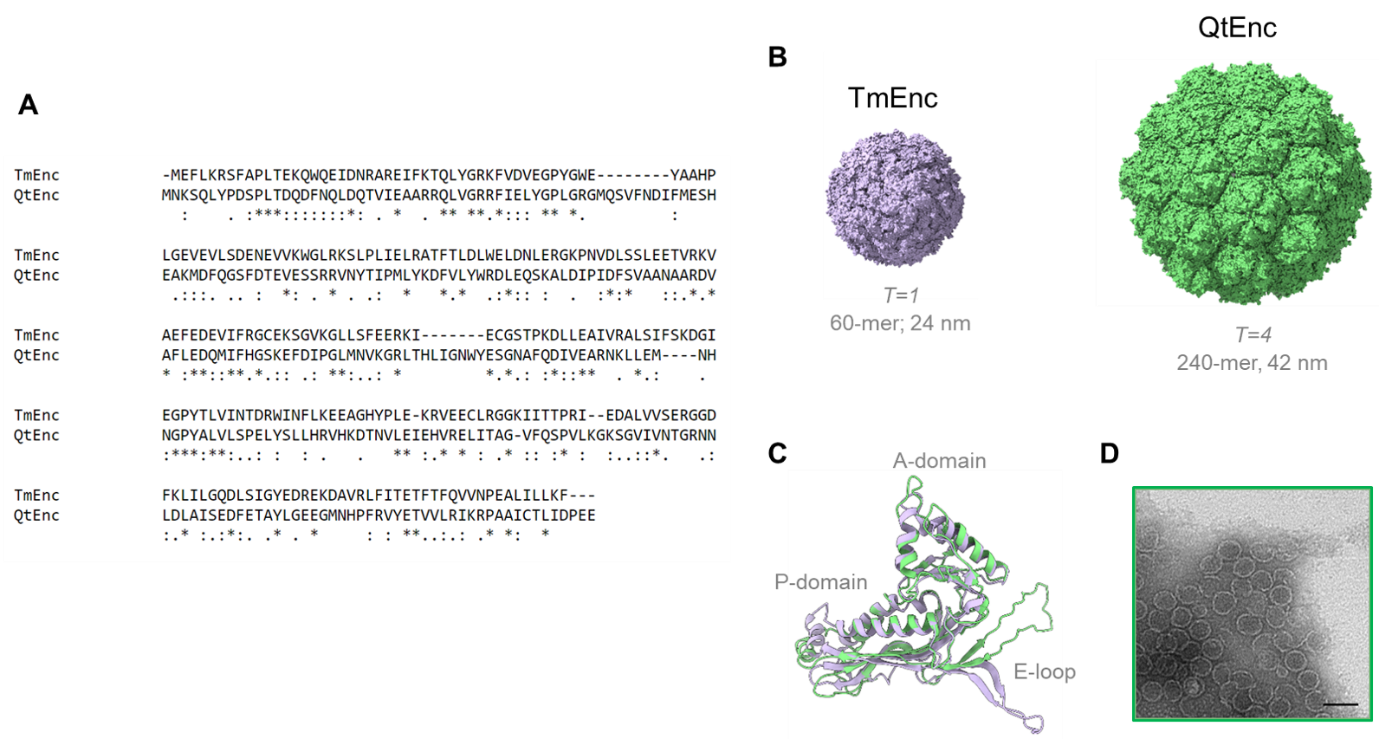

**Figure S5.** **The protein sequence and structure of *T*=1 TmEnc differs from *T*=4 QtEnc. (a)** Protein sequence alignment of TmEnc (PDB: 7KQ5) and QtEnc (PDB: 6NJ8) subunits indicates only ~22% of the amino acid residues are conserved between the two encapsulins. Annotations: (*****) fully conserved residue; (**:**) conservation between residues of highly similar properties; and (**.**) conservation between residues of weakly similar properties. Alignments performed in Clustal Omega.[1] **(b)** Relative size and structures of assembled TmEnc (*T*=1; 60-mer; 12 pentameric units; ~24 nm) and QtEnc (*T*=4; 240-mer; 12 pentameric and 30 hexameric units; ~42 nm).[2] **(c)** Structural overlay of the TmEnc (purple) and QtEnc (green) subunits. Like all other characterised encapsulins, both subunits possess a viral HK97-fold that enables their self-assembly into macromolecular nanocages.[3] However, structural differences are evident in their A-domain, P-domain, and especially their E-loop regions. All molecular graphics created with UCSF ChimeraX.[4] **(d)** TEM image of purified QtEnc nanocages used in this study (Scale bar = 50 nm).

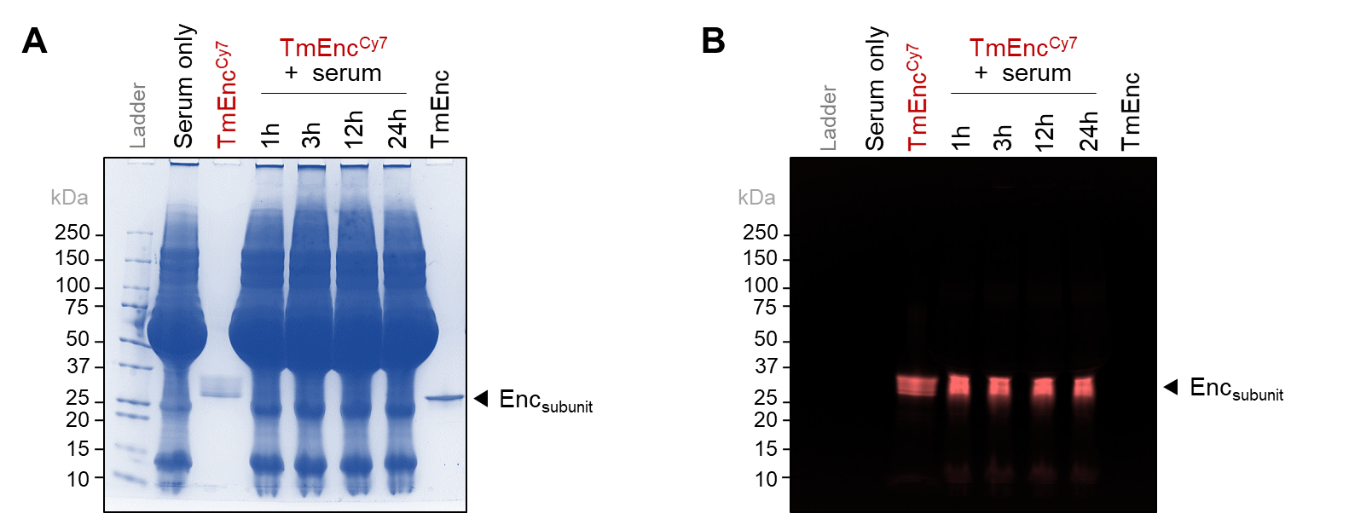

**Figure S6.** **Stability of dye-labelled TmEnc^Cy7^ in mouse serum.** SDS-PAGE analysis of TmEnc^Cy7^ incubated with mouse serum at 37°C for up to 24 h. **(a)** Coomassie stained proteins; and **(b)** In-gel Cy7 fluorescence.

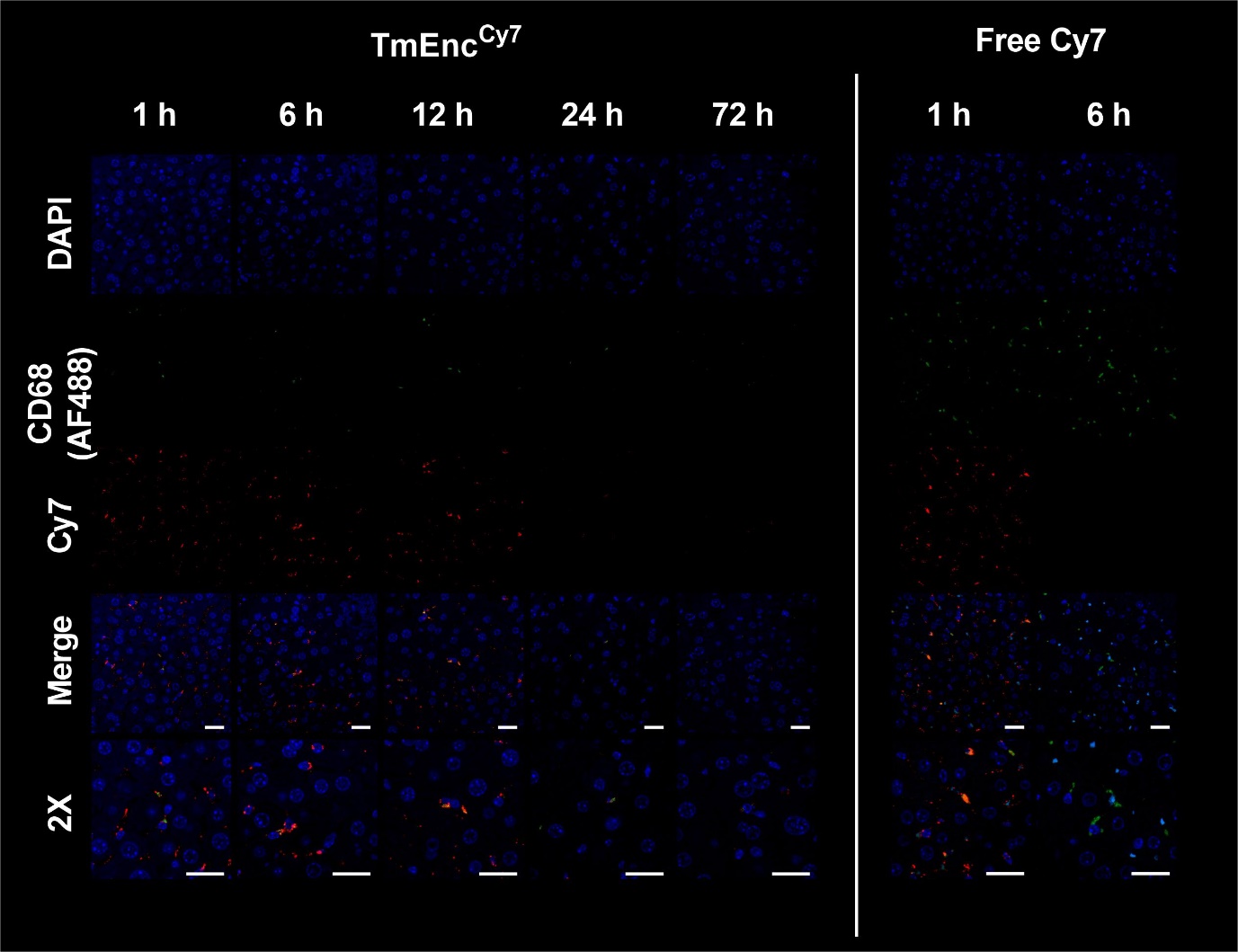

**Figure S7. Visualisation of TmEnc^Cy7^ uptake by liver cells.** Liver tissue sections embedded in paraffin were subjected to staining with a macrophage marker, CD68, and examined using confocal microscopy. 1 h after administration, TmEnc^Cy7^ nanocages (red) were observed in liver sinusoids, while Kupffer cells (green), the resident macrophages of the liver, internalized TmEnc^Cy7^ by 6 h post-injection. Some free Cy7 dye was diffusely distributed within the liver and was also taken up by Kupffer cells after 1 h; however, it became undetectable by 6 h (Scale bars = 25 µm).
